# The targeted deletion of genes responsible for expression of the Mth60 fimbriae leads to loss of cell-cell connections in *M. thermautotrophicus* ΔH

**DOI:** 10.1101/2022.05.13.491833

**Authors:** Christian Fink, Gines Martinez-Cano, Jeremiah Shuster, Aurora Panzera, Largus T. Angenent, Bastian Molitor

## Abstract

This study was continued by the Environmental Biotechnology Group of the University of Tübingen *in memoriam* to Reinhard Wirth, who initiated the work on Mth60 fimbriae at the University of Regensburg.

Growth in biofilms or biofilm-like structures is the prevailing lifestyle for most microbes in nature. The first step to initiate biofilms is the adherence of microbes to biotic and abiotic surfaces. Therefore, it is important to elucidate the initial step of biofilm formation, which is generally established through cell-surface structures (*i.e*., cell appendages), such as fimbriae or pili, that adhere to surfaces. The Mth60 fimbriae of *Methanothermobacter thermautotrophicus* ΔH are one of only few known archaeal cell appendages that do not assemble *via* the type-IV assembly mechanism. Here, we report the constitutive expression of Mth60 fimbriae-encoding genes from a shuttle-vector construct, as well as the deletion of the Mth60 fimbriae-encoding genes from the genomic DNA of *M. thermautotrophicus* ΔH. We expanded our system for genetic modification of *M. thermautotrophicus* ΔH by an allelic-exchange method. While overexpression of the respective genes resulted in an increase of the Mth60 fimbriae, deletion of the Mth60 fimbriae-encoding genes led to a loss of Mth60 fimbriae in planktonic cells of *M. thermautotrophicus* ΔH. This either increased or decreased number of Mth60 fimbriae correlated with a significant increase or decrease of biotic cell-cell connections in the respective *M. thermautotrophicus* ΔH strains compared to the wild-type strain.

**Originality-Significance Statement:** *Methanothermobacter* spp. have been studied for the biochemistry of hydrogenotrophic methanogenesis for many years. However, due to the lack of genetic tools, the detailed investigation of certain aspects, such as regulatory processes, was not possible. Here, we amend our genetic toolbox for *M. thermautotrophicus* ΔH with an allelic exchange method. We report the deletion of genes that encode for the Mth60 fimbriae. Our findings provide a first insight into the regulation of the expression of these genes and reveal a role of the Mth60 fimbriae in the formation of cell-cell connections of *M. thermautotrophicus* ΔH.

## Introduction

Microbial biofilm formation, maintenance, and dispersion in various habitats has been investigated in numerous studies ^1-3^. For bacteria, a plethora of studies elucidated biofilm formation, especially for clinically relevant pathogenic species ^4-7^. However, knowledge about archaeal biofilm formation still remains in an early stage ^8,9^. Archaea are found in extreme habitats with respect to pH, temperature, or salinity, as well as in moderate conditions such as sea water, the human gut, and rice paddy fields. Thus, there must have been several ways evolved to colonize habitats with different environmental properties ^10^. In general, formation of a biofilm in new habitats is established through cell-surface molecules and structures, which enable microbes to attach, and therefore adhere to a surface in the respective habitat ^6,11^. One possibility is adherence *via* extracellular polymeric substances ^12^. However, this was more frequently described to be important in a later stage of colonization and not for the initial attachment ^3,12^. For archaea, this initial attachment to surfaces mostly relies on archaeal cell appendages, such as archaella, pili, fimbriae, and other specialized archaeal cell appendages ^13,14^. In general, archaella differ from all other cell appendages in two ways: **1)** the diameter, which is 10-15 nm compared to ∼5 nm for fimbriae and pili; and **2)** the ability to rotate, and therefore enable directed motility of the microbe ^13^. It was shown that several archaella structures allow for adherence to surfaces *via* adhesins on the archaella tip ^14,15^. Archaella and the majority of cell appendages that have been described for archaea so far, are assembled *via* the type-IV assembly mechanism ^16,17^. However, some archaeal cell appendages assemble by mechanisms that are different from the type-IV assembly mechanism such as the hami from *Altiarchaeum hamiconnexum* and the Mth60 fimbriae from *M. thermautotrophicus* ΔH ^18,19^. These archaeal cell appendages do not enable the microbe for motility but for adherence to abiotic surfaces and biotic adherence between microbes.

Here, we focused on the Mth60 fimbriae from *M. thermautotrophicus* ΔH. The Mth60 fimbriae were first described by Doddema, *et al*. ^20^. They differ from archaella by their diameter and a length of up to 5 µm ^13,20^. Planktonic wild-type *M. thermautotrophicus* ΔH cells contain between one to three Mth60 fimbriae. In contrast, cells that are adhered to surfaces were found to contain significantly higher numbers of Mth60 fimbriae per microbial cell ^19^. *M. thermautotrophicus* ΔH was shown to adhere to several distinct surfaces, such as glass, carbon coated gold, copper grids, and silicium wavers, *via* the Mth60 fimbriae ^19^. Additional to abiotic surfaces, also biotic cell-cell connections with surface adhered *M. thermautotrophicus* ΔH have been demonstrated ^19^.

The Mth60 fimbriae mainly consist of the major fimbrin protein Mth60, which is eponymous for the Mth60 fimbriae. The corresponding gene, *mth60*, is transcribed in two transcriptional units (*i.e*., operons), *mth58-mth60* and *mth60-mth61* (MTH_RS00275-MTH_RS00285, MTH_RS00285-MTH_RS00290). Therefore, the level of transcription of *mth60* is largely elevated compared to the other genes *mth58, mth59*, and *mth61* in the two operons ^21^. Recombinant Mth60 protein, produced in *Escherichia coli*, led to auto-assembly of filamentous fimbriae structures when incubated at 65°C in *M. thermautotrophicus* ΔH growth medium ^19,21,22^. This auto-assembly feature of recombinant Mth60 protein was patented for a potential application as heat-induced glue through solidification of the Mth60 protein at elevated temperatures ^22^. Furthermore, the auto-assembly feature indicated an extraordinary assembly mechanism of Mth60 fimbriae compared to the type-IV assembly mechanism that was described for the majority of cell appendages in archaea ^13^. The function of *mth58, mth59*, and *mth61*, which are the three genes that are co-transcribed with *mth60*, remain largely unknown. Auto-assembly tests of Mth59 together with Mth60 failed in assembling filamentous structures. However, additional bioinformatics modelling of the Mth59 protein structure indicated a potential chaperone function of Mth59 for Mth60 ^21^.

To further investigate the relevance of Mth60 fimbriae for biotic cell-cell connections ^19^, we expanded the genetic tool-box for *M. thermautotrophicus* ΔH ^23^ with suicide vectors for targeted gene deletion. This enabled us to delete the Mth60 fimbriae-encoding operons (*mth58*-*mth60* and *mth60*-*mth61*) from the genomic DNA of *M. thermautotrophicus* ΔH by an allelic-exchange method. We further generated a strain of *M. thermautotrophicus* ΔH that contained a shuttle-vector construct for the constitutive expression of the Mth60 fimbriae-encoding operons. We observed varying phenotypes and significantly different numbers of Mth60 fimbriae per microbe for the different strains, and thus we were able to elucidate the intraspecies adherence ability of *M. thermautotrophicus* ΔH.

## Materials and Methods

### Microbial strains, media, and cultivation conditions

For cloning/gene manipulation and DNA transfer into *Methanothermobacter thermautotrophicus* ΔH we utilized the *Escherichia coli* strains NEB stable (New England Biolabs, Frankfurt/Main, Germany) and S17-1 ^24^, respectively. We cultivated *E. coli* in LB medium that contained: sodium chloride, 10 g/L; yeast extract, 5 g/L; tryptone, 10 g/L. For solidified LB media plates, we added 1.5 weight% of Kobe I Agar (Carl Roth, Karlsruhe, Germany). We supplemented LB medium with antibiotic substances for complementary antibiotic resistance genes on plasmids and integrated into genomic DNA of *E. coli* S17-1, with chloramphenicol, 30 µg/mL (plasmids) and trimethoprim, 10 µg/ml (*E. coli* S17-1). We incubated all *E. coli* cultures at 37°C. We incubated solidified media plates upside down in a static incubator, and liquid cultures in a shaker incubator with rotation (150 rpm).

For genetic modification and phenotypical analysis, we purchased *M. thermautotrophicus* ΔH (DSM 1053) from the DSMZ (Braunschweig, Germany). We cultivated *M. thermautotrophicus* ΔH in mineral medium that contained: sodium chloride, 0.45 g/L; sodium hydrogen carbonate, 6.00 g/L; di-potassium hydrogen phosphate, 0.17 g/L; potassium di-hydrogen phosphate, 0.23 g/L; ammonium chloride, 0.19 g/L; magnesium chloride hexahydrate, 0.08 g/L; calcium chloride dihydrate, 0.06 g/L; ammonium nickel sulfate, 1 mL (0.2 weight%); iron(II)chloride pentahydrate, 1 ml (0.2 weight%); resazurin indicator solution, 4 mL (0.025 weight%); and trace element solution, 1 mL (10-fold as stated in Balch and Wolfe ^25^). All chemicals were *per analysis* (*p.a*.) grade. We did not add vitamins. For solidified media plates, we added 1.5 weight% Bacto™ agar (BD Life Science, Berkshire, UK) *prior* to autoclaving. Neomycin sodium salt was supplemented for cultivation of genetically modified *M. thermautotrophicus* ΔH strains with concentrations of 250 µg/mL in liquid mineral media and 100 µg/mL in solidified media plates at 60°C, respectively.

We performed media preparation on the basis of anaerobic techniques as stated in Balch and Wolfe ^25^ with the modifications described in Fink, *et al*. ^23^. In brief, the composed media was sparged with N_2_/CO_2_ (80/20 volume%). Afterwards, for liquid media, we reduced the media with 0.5 g/L cysteine hydrochloride, dispensed the medium in serum bottles with a liquid/headspace ratio of 20 mL/80 mL (v/v) in an anaerobic chamber with a 100% N_2_ atmosphere (UniLab Pro Eco, MBraun, Garching, Germany), and performed a gas exchange to H_2_/CO_2_ (80/20 volume%). For solidified media plates, in addition, we added 0.3 g/L sodium sulfide monohydrate and dispensed 80 mL into 100-mL serum bottles. We exchanged the gas phase to N_2_/CO_2_ (80/20 volume%), and boiled the media to liquefy it directly *prior* to use. The amount of 80 mL per serum bottle was sufficient for 3-4 solidified media plates. We dried the plates for two hours in the anaerobic chamber. Afterwards, *M. thermautotrophicus* ΔH cell suspension could be plated on the surface ^23^. We incubated solidified media plates in pressurized stainless-steel jars (Raff und Grund, Freiberg, Germany) with an H_2_/CO_2_ (80/20 volume%) headspace at 60°C. *M. thermautotrophicus* ΔH in liquid culture was incubated rotating with 150 rpm at 60°C.

### Molecular cloning and vector construction

All primers, gBlocks (IDT, Coralville, IA, USA), and plasmids from this study are given in **Supplementary Table S1-S3**. We generated the template genomic DNA from *M. thermautotrophicus* ΔH for PCR amplification by using a gDNA extraction kit (Macherey+Nagel, Düren, Germany) with slight modifications. Instead of bead beating, we vortexed the microbe suspension for 1 min with 4-s intervals and eluted genomic DNA in 50 µL water instead of 100 µL elution buffer. We performed PCR amplification of vector and insert fragments with Q5 high-fidelity polymerase (New England Biolabs, Frankfurt/Main, Germany), followed by *Dpn*I restriction enzyme digest when necessary. We purified all PCR products *via* a PCR purification kit (Qiagen, Hilden, Germany), and extracted vector DNA from *E. coli via* QIAprep Spin Miniprep Kit (Qiagen, Hilden, Germany) *prior* to restriction enzyme digestion. Afterwards, we performed the restriction enzyme digestion and fragment ligation according to the manufacturer’s manual. We purchased all enzymes from New England Biolabs, Frankfurt/Main, Germany. We (re)transformed *E. coli* with vector constructs by chemical transformation following a standard heat-shock protocol ^26^. We confirmed all plasmids and vectors by Sanger sequencing performed at Genewiz (Azenta Life Sciences, Griesheim, Germany).

We generated a shuttle-vector construct for constitutive expression of the Mth60-fimbriae operons. For this, we PCR amplified the genes *mth58-mth61* (MTH_RS00275-MTH_RS00290) without the putative native promoter region upstream of *mth61* with primers Res_CF8 + Res_CF10 (**Supplementary Table S1**). Afterwards, we fused this PCR product with a gBlock containing the P_*hmtB*_ promoter *via* overlap extension PCR. This PCR amplicon contained the modular restriction sites *Pac*I and *Asc*I and could, thus, be fused after restriction enzyme digestion to the *Pac*I and *Asc*I digested pMVS1111A:P_*hmtB*_-*bgaB*, resulting in pMVS1111A:P_*hmtB*_-*mth58-61* ***(Supplementary* Table S3**).

To generate suitable suicide-vector constructs for genome integration of a thermostable neomycin resistance gene at the Mth60 fimbriae-encoding operon site, we deployed a three-step cloning strategy (**Supplementary Methods**). In brief, first the *E. coli* backbone was fused with 1-kb upstream and second with the downstream homologous flanking regions of the Mth60 fimbriae-encoding operons with a construct containing the neomycin resistance (Neo^r^) and P_*mcrB*(*M v*.)_ as non-functional spacer in between. In a third step, we exchanged the spacer flanked by *Fse*I and *Asc*I as modular restriction enzyme recognition sites toward the functional selectable marker module with P_synth_ in further suicide-vector constructs ^23,27^.

### Transformation of *M. thermautotrophicus* ΔH

We performed transformation of *M. thermautotrophicus* ΔH with an interdomain conjugation protocol for DNA transfer from *E. coli* S17-1 to *M. thermautotrophicus* ΔH as described in Fink, *et al*. ^23^. Summarized in short, we centrifuged 10 mL of stationary *E. coli* S17-1 that contained the shuttle- or suicide-vector construct at 3700 rpm for 10 min at room temperature (Centrifuge 5920 R, rotor S-4×1000, Eppendorf, Hamburg, Germany). We mixed the *E. coli* S17-1 cell pellet with a cell pellet from 8 mL of a stationary *M. thermautotrophicus* ΔH culture that we stepwise harvested inside the anaerobic chamber at 12500 rpm for 4 min at room temperature (MySPIN™ 12 Mini Centrifuge, Thermo Scientific Waltham MA, USA). Afterwards, the cell suspension of *E. coli* and *M. thermautotrophicus* ΔH was spot-mated on a solidified medium plate containing 50 volume% LB medium without sodium chloride and 50 volume% mineral medium. After the cell suspension was completely absorbed, the plate was incubated for 24 h at 37°C in a pressurized stainless-steel jar. The incubated spot-mated cells were washed-off the plate and transferred into sterile anaerobic mineral medium and incubated for 4 h at 60°C for recovery, expression of the neomycin resistance gene, and counterselection against *E. coli*. After the recovery, the *M. thermautotrophicus* ΔH mutants were enriched in 250 µg/mL neomycin-containing selective liquid mineral medium. The stationary grown enrichment culture was spread-plated on selective solidified medium plates and individual clonal populations were subjected to further analysis *via* PCR.

### Confirmation of *M. thermautotrophicus* ΔH mutant strains *via* PCR analysis

For screening purposes, we resuspended an individual clonal population in 50 µL of nuclease-free water or used 0.1 mL of *M. thermautotrophicus* ΔH culture directly and boiled the suspension at 100°C for 12 min *prior* to using 1 µL of suspension for PCR analysis. Final analysis was performed with 1 µL of genomic DNA extractions of respective *M. thermautotrophicus* ΔH mutant strains as template DNA for 10 µL PCR reaction mixes. PCR analysis was performed using Phire plant PCR master mix (Thermo Scientific, Waltham MA, USA). The denaturation and annealing times were increased to 20 sec and to 10 sec, respectively. A total of 30 cycles were performed for all analyses. We observed false positive PCR signals for shuttle-vector DNA and suicide-vector constructs due to plasmid DNA carry-over from *E. coli* for up to two transfers after the non-selective liquid recovery step. From the third transfer on, plasmid DNA from *E. coli* was not detectable anymore in any of our experiments.

### Immuno-fluorescence staining

For immuno-fluorescence staining analysis, we placed 20 µL of late exponential *M. thermautotrophicus* ΔH culture on a poly-L-lysine coated glass slide (VWR, Darmstadt, Germany). After allowing cells to settle onto the glass slide for 20 min, we washed the glass slide three times, for 5 min each, with phosphate-buffered saline (PBS, pH 7.4). Afterwards, we applied the anti-Mth60-fimbriae antibody (1:2000 diluted; rabbit) ^19^ in PBS, containing 0.3 weight% BSA (Carl Roth, Karlsruhe, Germany) and incubated for 2 h. We washed the sample three times, for 5 min each, with PBS (pH 7.4). Then, we applied a goat anti-rabbit IgG (Thermo Fisher Scientific, Waltham (MA), USA) cross-adsorbed secondary antibody with Alexa Fluor 488 (1:2000 diluted) in PBS, containing 0.3 weight% BSA, and incubated for 1 h. To reduce the background, the incubation was followed by three additional washing steps with PBS (pH 7.4). After the sample was almost dry, we applied 10 µL of Invitrogen™ ProLong™ Gold Antifade Mountant with DAPI on the sample and covered with a cover glass. Prolong Gold Antifade Mountant was allowed to solidify at 4°C for 24 h *prior* to Airyscan imaging analysis.

We performed Airyscan imaging at the Max Planck Institute for Biology Tübingen BioOptics Facility using a laser scanning inverted confocal microscope (Zeiss LSM 780; Carl Zeiss AG, Oberkochen, Germany) with a 63X oil/1.4NA oil-immersion objective. We used the diode laser line 405 nm for the excitation of DAPI, while using the 488 nm Argon laser line for the excitation of Alexa Fluor 488-conjugated antibody.

For the Airyscan images, we used the add-on Airyscan detection unit (Carl Zeiss AG), set to super resolution mode (SR). For each area we acquired a z-stack of images and processed the obtained data set first through the Airyscan software, which operates a 3D deconvolution on top of the pixel reassignment ^28^, followed then by maximum intensity projection along the z axis, to easily visualize the collected information on a single plane.

### Scanning electron microscopy

For scanning electron microscopy (SEM) analysis, we coated glow-discharged glass slides with 30 μL of 0.1% poly-L-Lysine (PLANO, Wetzlar, item number 18026) and dried them in a 60°C incubator oven for 1 h. In a fumehood, we placed a prepared glass slide (coated-side up) at the bottom of each well of a 24-well plate. We separately added a 100-µL aliquot of each strain (*i.e*., late exponential culture of *M. thermautotrophicus* ΔH wild-type, constitutive expression, and deletion strains) to a well and cells were allowed to settle onto the glass slide. After 20 min of incubation, we removed the supernatant and added 100-µL PBS. Electron microscopy grade glutaraldehyde (25%, PLANO, Agar Scientific, item number R1011) was added to each sample to obtain an overall 2.5 volume%. Afterwards, we covered and incubated the 24-well plate at room temperature (ca. 21°C) for 1 h, allowing for fixation and for cells to settle onto the slides. After incubation, we removed the supernatant from each well and rinsed the sample-bearing slides by adding deionized water to each well and incubating for 10 min to remove material that did not attach to the glass slide. This rinsing procedure was repeated by removing the supernatant and adding fresh deionized water. After the final rinse, we dehydrated the samples using a graded ethanol series: 25, 50, 75 volume% ethanol (15 min incubation at each concentration), and three times 100 volume% ethanol (30 min incubations). After the last ethanol dehydration, we added 100 volume% hexamethyldisilazane (HMDS) to each sample so that each well contained a 50/50 volume% ratio of HMDS/100 volume% ethanol. Then, we covered the plate and allowed to incubate for 30 min. After incubation, we removed the HMDS/ethanol solution and added 100 volume% HMDS to each well. We left the plate lid partially open to allow airdrying to occur overnight. The sample-bearing glass slides were adhered to aluminum stubs using carbon adhesive tabs (PLANO, Wetzlar, item numbers G301 & G3347) and coated with ca. 8 nm of platinum using a BAL-TEC™ SCD 005 sputter coater. We performed the structural characterization of *M. thermautotrophicus* ΔH strains using a Zeiss Crossbeam 550L Focused Ion Bean (FIB) – Scanning Electron Microscope (Oberkochen, Germany), operating with an acceleration voltage of 2 kV. We took all micrographs using secondary electron (SE) mode.

### Phase-contrast microscopy analysis

We placed 10 µL of untreated late exponential (OD_600_=0.28) *M. thermautotrophicus* ΔH cultures (each of wild-type, constitutive expression, and deletion strain) on microscopy slides and added cover slips (N=3 for each strain). After 10 min of incubation at room temperature, 30 pictures (10 pictures of each replicate) for all three *M. thermautotrophicus* ΔH strains were taken. For this, we chose random vision fields at 100-fold magnification and phase contrast 3 with System Microscope BX41TF (Olympus, Shinjuku, Japan; equipped with a U-TV0.5XC-3 camera).

In all 30 pictures for each *M. thermautotrophicus* ΔH strain, we counted the total number of microbes and the number of microbes, which showed connection to another microbe. Afterwards, we calculated the ratio between the total number and number of connected microbes using R ^29,30^.

## Results

### Suicide-vector constructs allow site-specific deletion of Mth60 fimbriae-encoding operons in *M. thermautotrophicus* ΔH

We chose suicide-vector constructs to substitute *mth58-mth61* (*i.e*., the Mth60 fimbriae-encoding operons MTH_RS00275-MTH_RS00290) with a positive selectable marker. Therefore, we created suicide-vector constructs with ∼1-kb homologous flanking regions upstream and downstream of the Mth60 fimbriae-encoding operons (**Figure 1**). We placed unique restriction enzyme-recognition sites at the interfaces between the homologous flanking regions, which do rarely/do not occur in the genome of *M. thermautotrophicus* ΔH. While *Sal*I and *Not*I occur 137 and 7 times on the genome of *M. thermautotrophicus* ΔH, respectively, *Asc*I and *Fse*I are not present at all. By using these restriction enzyme-recognition sites, we ensured that exchangeability with virtually all homologous flanking regions for *M. thermautotrophicus* ΔH genes is possible. This modularity facilitates the generation of future suicide-vector constructs. We implemented the restriction enzyme-recognition sites *Asc*I and *Fse*I at the selectable-marker interfaces. With that, it is also possible to directly implement selectable markers from the pMVS shuttle-vector design for *M. thermautotrophicus* ΔH into suicide vector constructs ^23^. As selectable marker, we used the thermostable neomycin resistance gene under the control of the P_Synth_ promoter ^23,31,32^. The commonly used T_*mcr*(*M. v.*)_ sequence served as terminator ^33^.

**Figure 1.**
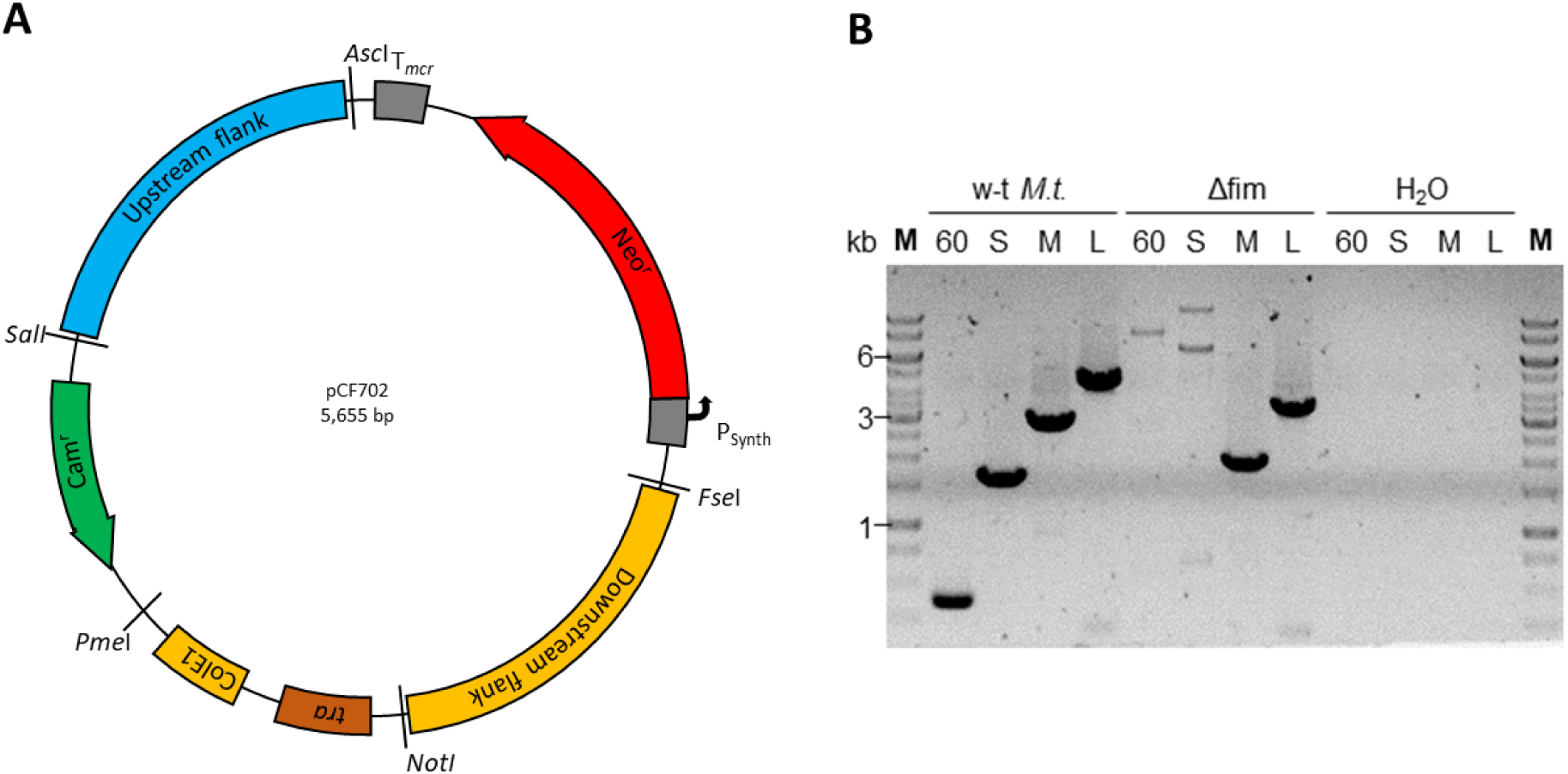
Plasmid map of pCF702, a suicide-vector construct for deletion of the Mth60 fimbriae-encoding operons, (**A**) and the agarose gel of a corresponding PCR analysis (**B**). **A)** The suicide-vector construct pCF702 consists of five exchangeable modules flanked by unique restriction enzymes-recognition sites as stated in parentheses below. The origin of replication **ColEI** for *E. coli* including a ***tra*** region for mobilization during conjugation (*Not*I, *Pme*I), the antibiotic resistance against chloramphenicol (**Cam**^R^) as selectable marker for *E. coli* (*Pme*I, *Sal*I), 1-kb upstream (*Sal*I, *Asc*I) and downstream homologous regions (*Fse*I, *Not*I) for homologous recombination in *M. thermautotrophicus* ΔH. In between the homologous flanking regions, a thermostable neomycin resistance gene (**Neo**^R^) with constitutive promoter **P**_Synth_ and terminator **T**_*mcr*_ as selectable marker for *M. thermautotrophicus* ΔH is located (*Asc*I, *Fse*I). **B)** PCR analysis with four primer combinations to confirm the Mth60-fimbriae operon deletion. Two primer combinations amplify a fragment inside the Mth60 fimbriae-encoding operons (**60, S**), two primer combinations amplify a fragment outside-outside the Mth60 fimbriae-encoding operons (**M, L**). These combinations result in amplified fragments of reduced lengths since the Mth60 fimbriae-encoding operons (2.8 kb) were substituted with Neo^R^ (1.2 kb).

We performed the transformation of *M. thermautotrophicus* ΔH with the suicide-vector construct as described before ^23^. However, for plating of *M. thermautotrophicus* ΔH deletion mutants we applied 100 µg/mL of neomycin instead of 250 µg/mL, which we used for liquid selective media. This ensured the generation of individual clonal populations on selective solidified media plates because no clonal population appeared on plates with the higher antibiotic concentration. The generation of a clean *M. thermautotrophicus* ΔH strain with a deletion of the Mth60 fimbriae-encoding operons was challenging, and we continuously found wild-type signals in the PCR analysis in addition to the correct signal for double-homologous recombination events (**Supplementary Figure S2**). Nanopore sequencing of one of these cultures with mixed PCR signals revealed the co-existence of single- and double-homologous recombination events of the suicide vector with genomic DNA of *M. thermautotrophicus* ΔH (*Supplementary Figure S4*), while wild-type *M. thermautotrophicus* ΔH nanopore sequencing reads did not align with the neomycin resistance gene (**Supplementary Figure S3**). After an additional screening step with four individual clonal populations, we were able to isolate a *M. thermautotrophicus* ΔH *Δmth58-61*::NeoR mutant without wild-type genomic DNA background (**Figure 1B**). We confirmed the absence of wild-type *mth58-61* with the help of two specific primer combinations. Furthermore, we determined the substitution of the Mth60 fimbriae-encoding operons with the neomycin selectable marker with two additional specific primer pairs. The latter primer combinations would result in two PCR fragments, when wild-type *M. thermautotrophicus* ΔH genomic DNA background was still present. Thus, the uniformity of the genotype, and therefore the purity of *M. thermautotrophicus* ΔH *Δmth58-61*::NeoR strain was confirmed (**Figure 1B**).

Additional to the Mth60 fimbriae deletion strain of *M. thermautotrophicus* ΔH, we generated a *M. thermautotrophicus* ΔH strain (*M. thermautotrophicus* ΔH pMVS1111A:P_*hmtB*_-*mth58-61*) that constitutively expressed the Mth60 fimbriae-encoding operons. For this, we exchanged the gene of interest module of the pMVS1111A:P_Synth_-*bgaB* shuttle vector with the *mth58-mth61* genes under the control of the P*_hmtB_* promoter, which substituted the putative promoter region that is located upstream of *mth61* (**Supplementary Figure S1A**)^23^. After transformation of wild-type *M. thermautotrophicus* ΔH with the shuttle-vector construct for constitutive expression, we confirmed the maintenance of the construct after three and four transfers of the culture with a specific primer combination for the origin of replication module (***Supplementary Figure S1B***).

### Constitutive expression of Mth60 fimbriae-encoding operons results in an increase, and deletion results in a loss of visualizable Mth60 fimbriae compared to wild-type *M. thermautotrophicus* ΔH

The *M. thermautotrophicus* ΔH mutant strains for constitutive expression and with the deletion of Mth60 fimbriae-encoding operons allowed us to compare the resulting phenotypes to wild-type *M. thermautotrophicus* ΔH and with each other. For analysis of the phenotypes, we chose two distinct microscopical approaches to visualize Mth60 fimbriae. First, we used immuno-fluorescence staining with confocal light microscopy, and second, scanning electron microscopy of native *M. thermautotrophicus* ΔH (mutant) strain samples.

For immuno-fluorescence staining, we applied an anti-Mth60-fimbriae antibody as the first antibody. This antibody was generated by Christina Sarbu from the University of Regensburg from a density gradient centrifugation fraction with a high content of Mth60 fimbriae ^21^. The anti-Mth60-fimbriae antibody does bind to Mth60 fimbriae. However, it was also shown to bind to other cell-membrane components. This resulted in immuno-fluorescence staining of the entire *M. thermautotrophicus* ΔH cell additionally to the Mth60 fimbriae (**Figure 2, larger field of view in Supplementary Figure S5**). We passively attached planktonic wild-type *M. thermautotrophicus* ΔH from liquid media on poly-lysine glass slides. After immuno-fluorescence staining, we found one to a few stained Mth60 fimbriae per planktonic wild-type *M. thermautotrophicus* ΔH cell, which provided us with the necessary proof-of-principle for the success of the immuno-staining procedure (**Figure 2A**). Similar numbers of Mth60 fimbriae for planktonic wild-type *M. thermautotrophicus* ΔH were also described in Thoma, *et al*. ^19^. Thus, we analyzed specimens of the *M. thermautotrophicus* ΔH pMVS1111A:P_*hmtB*_-*mth58-61* that constitutively expressed Mth60 fimbriae, and we found a number of Mth60 fimbriae per cell that largely exceeded those of stained wild-type *M. thermautotrophicus* ΔH cells (**Figure 2A+B**). On the other hand, we compared the *M. thermautotrophicus* ΔH *Δmth58-61*::NeoR strain to the wild-type strain with the same immuno-fluorescence staining procedure, and found that specimens of the *M. thermautotrophicus* ΔH *Δmth58-61*::NeoR strain contained the stained cell wall, but did not show any Mth60 fimbriae (**Figure 2C**). Additionally, we observed detached/solitary Mth60 fimbriae frequently in the constitutively expressing strain and low numbers for the wild-type strain, but never in the Mth60-fimbriae deletion strain.

**Figure 2.**
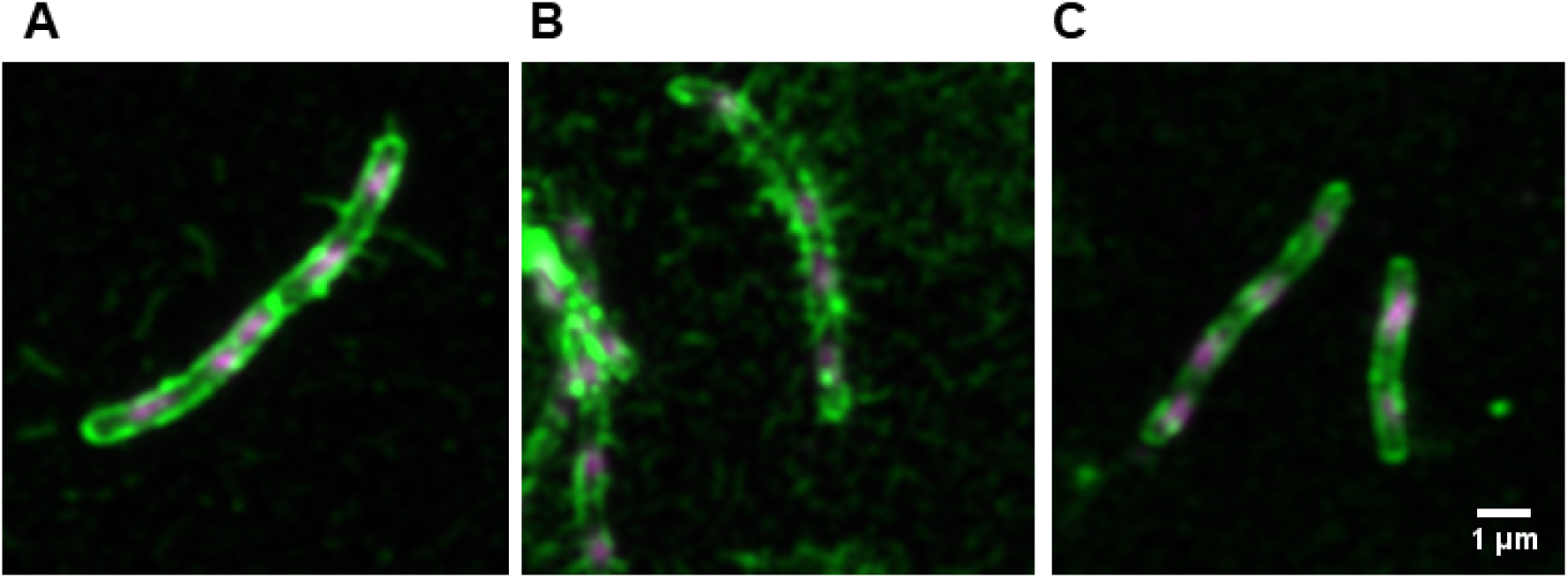
Two-channel maximum intensity z-projection of Airyscan processed z-stacks of immune-fluorescence-stained *M. thermautotrophicus* ΔH strains (A-C). Dapi staining is represented in magenta. The Alexafluor-488 conjugated antibody, which is attached to primary anti-Mth60-fimbriae antibody is depicted in green. The 1-µm scale bar represents the size for pictures A-C, because the magnification is the same. **A)** M. thermautotrophicus ΔH wild-type. **B)** *M. thermautotrophicus* ΔH containing a shuttle vector for constitutive expression of the Mth60 fimbriae-encoding operons. **C)** *M. thermautotrophicus* ΔH with a deletion of the Mth60 fimbriae-encoding operons.

The immuno-fluorescence staining enabled us to visualize varying numbers of Mth60 fimbriae for the different *M. thermautotrophicus* ΔH strains (wild-type, constitutive expression, and deletion strain). However, we aimed for another layer of evidence to confirm differences for the three strains, and decided to employ scanning electron microscopy. To maintain a state that is closest to the physiological conditions in the serum bottles, we passively attached the planktonic *M. thermautotrophicus* ΔH cells to poly-lysine coated SEM cover slips. This avoided centrifugation, and therefore potential disruption of Mth60 fimbriae. The results of scanning electron microscopical analysis of wild-type *M. thermautotrophicus* ΔH aligned with the observations from immuno-fluorescence staining and revealed in general one to a few stained Mth60 fimbriae per *M. thermautotrophicus* ΔH wild-type cell (**Figure 3A**). In the constitutive expression strain, the Mth60 fimbriae appeared to be more frequent than in specimens of wild-type *M. thermautotrophicus* ΔH (**Figure 3B**). These Mth60 fimbriae were visible as filaments that connect different cells with each other (**Figure 3A+B**). However, the difference did not appear as strong as indicated by the immuno-fluorescence staining procedure. We did not observe Mth60 fimbriae that connect microbes with each other in the Mth60-fimbriae deletion strain, such as we did for wild-type *M. thermautotrophicus* ΔH and the constitutive expression strain specimens (**Figure 3C**). In all specimens, including in the Mth60-fimbriae deletion strain, we found additional extracellular structures that were attached to cells, which: **1)** where round and condensed in shape; **2)** did not connect cells with each other; and **3)** did not resemble the filamentous structure of Mth60 fimbriae that we found only in wild-type *M. thermautotrophicus* ΔH and the constitutive expression strain.

**Figure 3.**
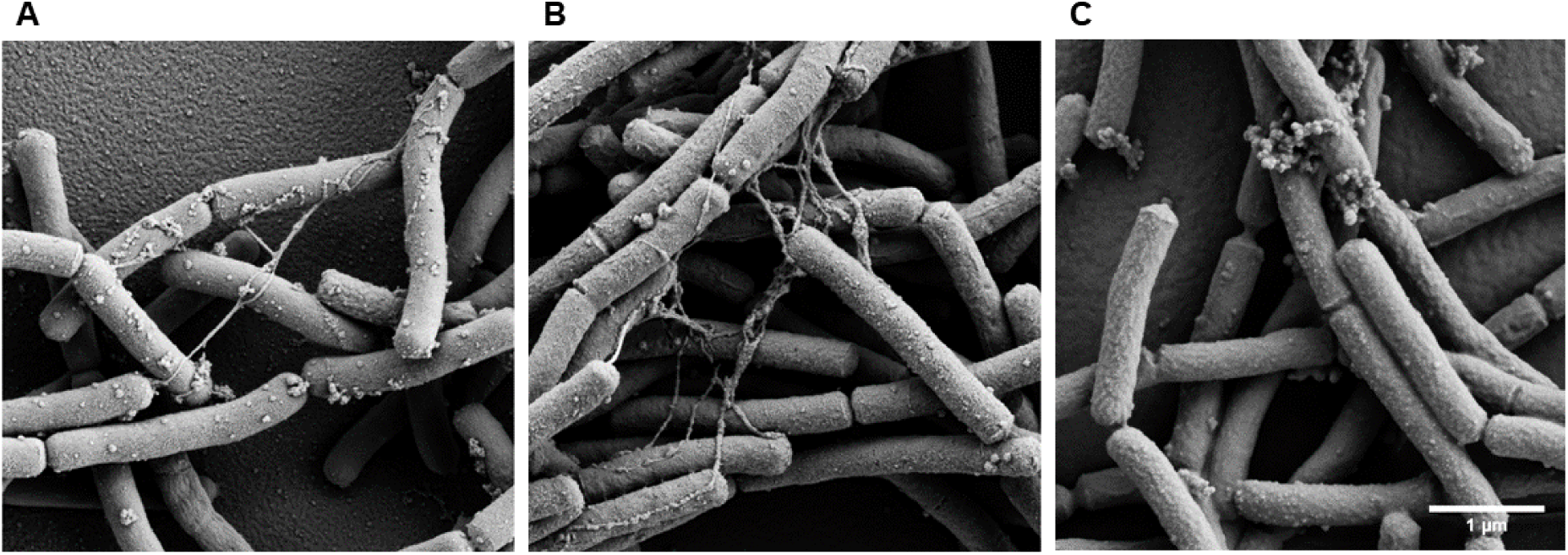
Scanning electron microscopy pictures of three untreated *M. thermautotrophicus* ΔH strains **(A-C)**. The scale bar is indicating 1 µm for A-C with a magnification of 10000x. **A)** *M. thermautotrophicus* ΔH wild-type. **B)** *M. thermautotrophicus* ΔH containing a shuttle vector for constitutive expression of the Mth60 fimbriae-encoding operons. **C)** *M. thermautotrophicus* ΔH with a deletion of the Mth60 fimbriae-encoding operons.

### The number of Mth60 fimbriae significantly influences intraspecies biotic interaction ability of *M. thermautotrophicus* ΔH

The experiments described above strongly indicated an increase of Mth60 fimbriae in the constitutive expression strain, as well as a loss of Mth60 fimbriae in the deletion strain, compared to wild-type *M. thermautotrophicus* ΔH. Thus, we hypothesized that these changes would be reflected in physiological differences between the strains. All three strains (wild-type, constitutive expression, and deletion strain) grew over night to a final optical density at 600 nm (OD600) of around 0.3. However, we observed a considerable difference between the strains with phase-contrast microscopy without further treatment in the early stationary growth phase. For wild-type *M. thermautotrophicus* ΔH, we found several cell clumps but also cells that remained planktonic (Figure 4A). For the constitutive expression strain, the ratio of cell clumps to planktonic cells shifted towards cell clumps (**Figure 4B**), while for the deletion strain it shifted towards planktonic cells (**Figure 4C**). This finding prompted us to develop a method to define significant differences in the number of cell clumps, and thus to determine a significant physiological difference between the three strains. Therefore, we collected a relevant number of phase-contrast microscopy pictures (n=10) from biological replicates (N=3) for each strain. Afterwards, we counted: **1)** the total number of microbes in each picture; and **2)** the number of microbes that had an immediate connection to another microbe. It was crucial to dilute the cells to the same optical density (OD_600_=∼0.28) to gather comparable results. The ratio of the number of total cells to connected cells resulted in significant differences in cell-cell connections for the three strains (**Figure 4D**). While in wild-type *M. thermautotrophicus* ΔH, 34.5±13% of the cells showed a connection to another cell, this was 52±15% in the constitutive expression strain. The Mth60-fimbriae deletion strain only showed a remaining 7.5±4.5% of cells that were connected to other cells.

**Figure 4.**
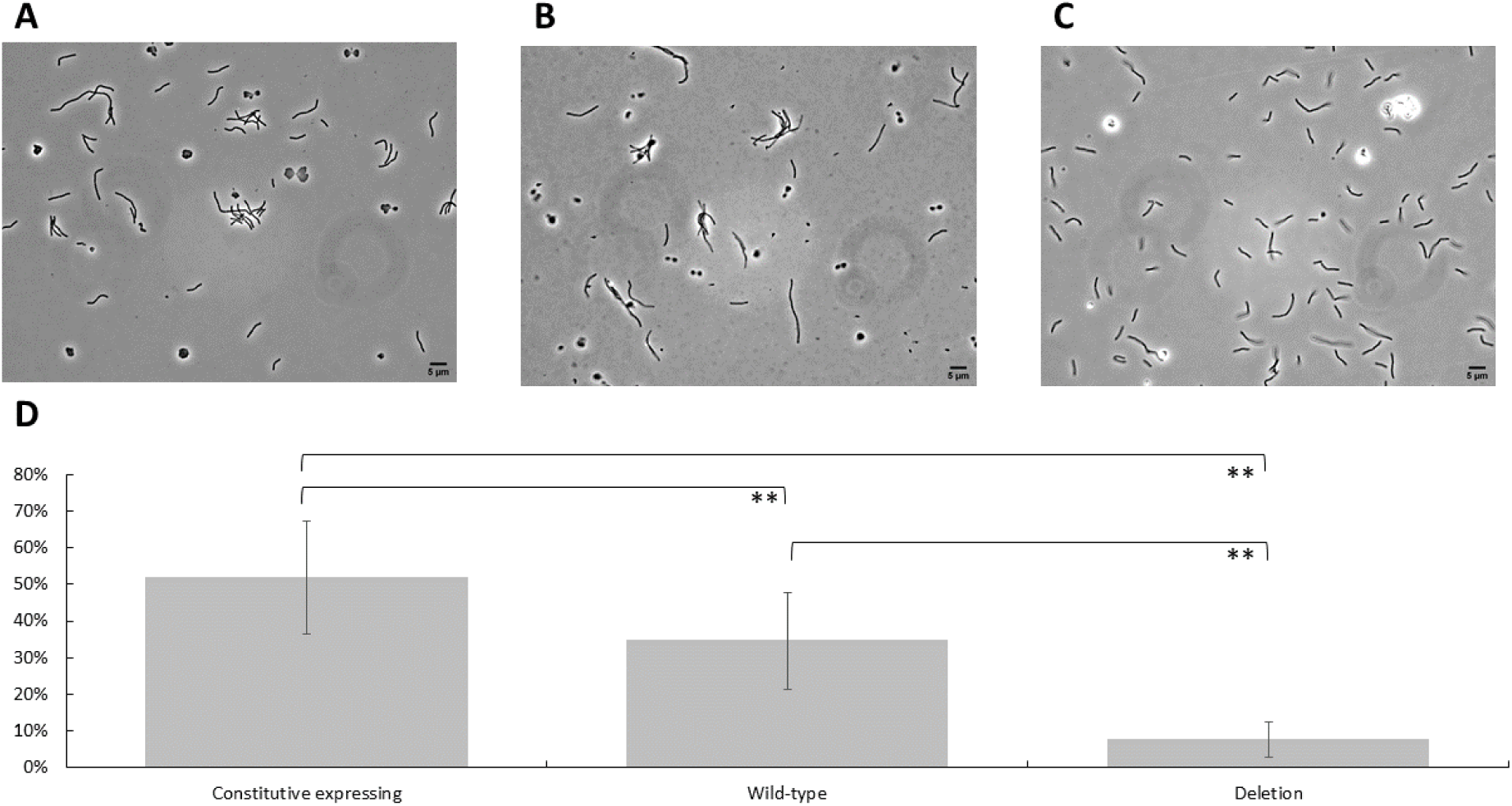
Phase-contrast microscopy pictures with magnification of 1000x of three *M. thermautotrophicus* ΔH strains **(A-C)** as representative pictures on which basis the analysis of the cell-cell connections was performed **(D). A)** Wild-type *M. thermautotrophicus* ΔH. **B)** Constitutive Mth60 fimbriae expressing *M. thermautotrophicus* ΔH. **C)** Mth60 fimbriae deletion strain of *M. thermautotrophicus* ΔH. D) Comparison of the three *M. thermautotrophicus* ΔH strains mentioned above regarding total number of microbial cells with the number of microbial cells connected to another cell in percent. Average (N=3, n=10) with error bars indicating standard deviation. Significance was tested with Student’s t-test (two-tailed): *, significant difference (P<0.05); **, highly significant difference (P<0.01).

## Discussion

In this study, we reported the implementation of suicide-vector constructs for homologous recombination in *M. thermautotrophicus* ΔH to generate site-specific gene deletion mutants *via* allelic exchange with a positive selectable marker. With our expanded genetic tools, we elucidated the positive and negative influence of constitutive expression and deletion of the Mth60 fimbriae-encoding operons on the *in-vivo* production of Mth60 fimbriae in *M. thermautotrophicus* ΔH. We demonstrated a correlation between the number of Mth60 fimbriae and the number of cell-cell connections with a constitutive Mth60-fimbriae expression strain, wild-type strain, and Mth60-fimbriae deletion strain of *M. thermautotrophicus* ΔH. We measured significantly lower numbers of cell-cell connections in *M. thermautotrophicus* ΔH strains with lower numbers of Mth60 fimbriae, and therefore demonstrated the importance of Mth60 fimbriae for the establishment of cell-cell connections, which is essential for initial biofilm formation.

The DNA-transfer protocol, and therefore the generation of deletion mutants of M. thermautotrophicus ΔH, was performed with the identical procedure as we had established for shuttle-vector constructs before ^23^. However, for the successful isolation of mutant strains, the concentration of neomycin as the antibiotic substance had to be lowered to 100 µg/mL instead of 250 µg/mL on solidified media plates when a genomic alteration was introduced. It is known that cells can adapt the copy number of plasmids in response to higher antibiotic-substance concentrations, which leads to higher resistance levels towards these antibiotic substances ^34^. It is further known that *M. thermautotrophicus* ΔH is always diploidic ^35^. Thus, we argue that the copy number of our shuttle vector is likely higher than two (as for the genome copies) or potentially can be increased with higher antibiotic-substance concentrations. This would explain higher neomycin resistance levels of shuttle-vector containing *M. thermautotrophicus* ΔH compared to genome-altered *M. thermautotrophicus* ΔH mutant strains.

During the procedure of isolating a clean *M. thermautotrophicus* ΔH strain with a deletion of the Mth60 fimbriae-encoding operons, PCR signals and Nanopore sequencing reads for wild-type *M. thermautotrophicus* ΔH, single-homologous recombined, and double-homologous recombined mutant strains were obtained from the same colony sample, even after two steps that included the isolation of an individual clonal population and the transfer to liquid growth medium (**Supplementary Figure S2**). One possible explanation is the diploid character of *M. thermautotrophicus* ΔH, which might result in residual wild-type or single-homologous recombined alleles on the second chromosome ^35^. This could result in a heterozygous culture of *M. thermautotrophicus* ΔH as it was shown to appear in heterozygous and many genome copies-containing *Methanococcus maripaludis* cultures ^36^. Another possible explanation is the characteristic of *M. thermautotrophicus* ΔH of forming multicellular filaments. This could result in different genotypes in one filament of multiple individual *M. thermautotrophicus* ΔH cells ^35,37^. These observations of various genotypical PCR signals make it difficult, but not impossible, to isolate clean deletion strains of *M. thermautotrophicus* ΔH (**Figure 1B**).

We performed immuno-fluorescence staining to visualize the Mth60 fimbriae with the Mth60-fimbriae deletion, the constitutive Mth60-fimbriae producing, and wild-type *M. thermautotrophicus* strains. The Mth60-fimbriae antibodies, that we used for immuno-fluorescence staining, were generated from a native Mth60-fimbriae preparation, which was purified through density gradient centrifugation. After our staining approach, we demonstrated that in addition to the Mth60 fimbriae also the entire cell wall was stained, which resulted in a staining of the entire cell (**Figure 2A-C**). One possible explanation is that cell-wall components were purified in the same fraction of the density gradient centrifugation, resulting in a mixture of the polyclonal antibodies against several antigens. Another explanation is that the Mth60 fimbriae antibody recognizes glycosylated epitopes of the major fimbrin Mth60 of the Mth60 fimbriae ^19^. In that case, the Mth60-fimbriae antibody might also bind glycosylated cell-wall components on the envelope of *M. thermautotrophicus* ΔH cells ^38^.

Thoma, *et al*. ^19^ mentioned a difference in the number of Mth60 fimbriae in planktonic *M. thermautotrophicus* ΔH cells *vs*. cells that were actively grown in the presence of a surface to which the cells adhered. While only 50% of planktonic cells contained few Mth60 fimbriae, cells that were adhered to surfaces contained large numbers of Mth60 fimbriae per microbial cell ^19^. This finding clearly indicated a regulation of the expression of the Mth60 fimbriae-encoding operons. When we exchanged the putatively regulated promoter to the constitutive P_*hmtB*_ promoter, fimbriae were identified in higher numbers for each planktonic *M. thermautotrophicus* ΔH cell (**Figure 2, 3**) ^23^. The regulatory mechanism of putative promoter regions of the Mth60 fimbriae-encoding operons, however, will need to be investigated further.

The Mth60-fimbriae deletion mutant of *M. thermautotrophicus* ΔH does not contain any Mth60 fimbriae (**Figure 2C, 3C**). This loss of Mth60 fimbriae did not influence the generation of individual multicellular filaments, however, the connections to other multicellular filaments was significantly reduced (**Figure 4**). From this, we concluded that Mth60 fimbriae are the only cell appendages of *M. thermautotrophicus* ΔH that are responsible for biotic intraspecies cell-cell connections under the conditions that we investigated. Furthermore, we argue that Mth60 fimbriae are not involved in the formation of multicellular filaments, as these filaments were present in all *M. thermautotrophicus* ΔH strains that we analyzed. It was shown that the addition of Mth60-fimbriae antibodies to surface-adhered *M. thermautotrophicus* ΔH cells led to detachment of the cells, potentially by blocking the Mth60 fimbriae adhesion mechanism ^19^. With the deletion of the Mth60-fimbriae operons, and therefore the loss of Mth60 fimbriae, we were now able to support these results on a genetic level by demonstrating reduced cell-cell connections *in vivo*.

We demonstrated that deletion of all four genes that are co-transcribed with *mth60*, including *mth60*, led to the loss of Mth60 fimbriae. In addition, we provided further evidence for the regulation of the Mth60 fimbriae-encoding operons. Based on these findings, the functions of the individual genes in the Mth60 fimbriae-encoding operons can be studied in further detail now. The putatively regulated promoters of the Mth60 fimbriae-encoding operons are the first step towards the identification of a sensory system in *M. thermautotrophicus* ΔH that allows adherence to biotic and abiotic surfaces for initial biofilm formation. The reduced ability to form cell-cell connections might have an impact on the rheology of a high-density microbial culture, and thus may affect the biotechnological applications with *M. thermautotrophicus*, such as for power-to-gas processes, in large-scale fermentation to convert carbon dioxide and hydrogen to renewable methane ^39^. Clearly, a possible effect of the rheology on parameters, such as mixing, gas solubility, and gas conversion efficiency, with the pili-deficient strain of *M. thermautotrophicus* ΔH will have to be addressed in future research.

## Acknowledgments

The authors thank Marco-Linus Ernst and Andreas Mark Enkerlin for their support with wet lab experiments. In addition, the authors thank the Archaea Centre of the University of Regensburg with Annett Bellack for the Mth60-fimbriae antibodies. The work was funded by the Alexander von Humboldt Foundation in the framework of the Alexander von Humboldt Professorship (L.T.A.) and the U.S. Office of Naval Research Global (ONRG, N62909-19-1-2076; L.T.A., B.M.). We thank the additional funding sources, which were the German Federal Ministry of Education and Research (MethanoPEP, 031B0851C, B.M.; ThermoSynCon, 031B0857D, L.T.A), the Deutsche Forschungsgemeinschaft (DFG, German Research Foundation) under Germany’s Excellence Strategy – EXC 2124 – 390838134 (L.T.A., B.M.), and the DFG (INST 37/1027-1 FUGG) for financial support provided for the acquisition of the cryogenic focused ion beam scanning electron microscope.

## Author Contributions

B.M. and L.T.A. initiated the work. C.F. and B.M. designed the experiments. J.S. and C.F. did the preparation and analysis of scanning electron microscopy. A.P. performed Airyscan microscopy and analysis. C.F. and G.M-C. performed laboratory experiments and analyzed the data. L.T.A. and B.M. supervised the project. C.F. wrote the manuscript, while all edited the paper and approved the final version.

## References

1 Rumbaugh, K. P. & Sauer, K. Biofilm dispersion. Nat Rev Microbiol 18, 571–586 (2020).

2 Hobley, L., Harkins, C., MacPhee, C. E. & Stanley-Wall, N. R. Giving structure to the biofilm matrix: An overview of individual strategies and emerging common themes. FEMS Microbiol Rev 39, 649–669 (2015).

3 Davey, M. E. & O’toole, G. A. Microbial biofilms: From ecology to molecular genetics. Microbiol Mol Biol Rev 64, 847–867 (2000).

4 Pelling, H. et al. Bacterial biofilm formation on indwelling urethral catheters. Lett Appl Microbiol 68, 277–293 (2019).

5 Jamal, M. et al. Bacterial biofilm and associated infections. Chin Med J 81, 7–11 (2018).

6 O’Toole, G., Kaplan, H. B. & Kolter, R. Biofilm formation as microbial development. Annu Rev Microbiol 54, 49–79 (2000).

7 Ciofu, O., Moser, C., Jensen, P. Ø. & Høiby, N. Tolerance and resistance of microbial biofilms. Nat Rev Microbiol, 1–15 (2022).

8 Wirth, R. Colonization of black smokers by hyperthermophilic microorganisms. Trends Microbiol 25, 92–99 (2017).

9 van Wolferen, M., Orell, A. & Albers, S.-V. Archaeal biofilm formation. Nat Rev Microbiol 16, 699–713 (2018).

10 Lyu, Z., Shao, N., Akinyemi, T. & Whitman, W. B. Methanogenesis. Curr Biol 28, R727–R732 (2018).

11 Palmer, J., Flint, S. & Brooks, J. Bacterial cell attachment, the beginning of a biofilm. J Ind Microbiol Biotechnol 34, 577–588 (2007).

12 Flemming, H.-C. & Wingender, J. The biofilm matrix. Nat Rev Microbiol 8, 623–633 (2010).

13 Jarrell, K. F., Ding, Y., Nair, D. B. & Siu, S. Surface appendages of archaea: Structure, function, genetics and assembly. Life 3, 86–117 (2013).

14 Chaudhury, P., Quax, T. E. F. & Albers, S. V. Versatile cell surface structures of archaea. Mol Microbiol 107, 298–311 (2018).

15 Näther, D. J., Rachel, R., Wanner, G. & Wirth, R. Flagella of Pyrococcus furiosus: Multifunctional organelles, made for swimming, adhesion to various surfaces, and cell-cell contacts. J Bacteriol 188, 6915–6923 (2006).

16 Pohlschroder, M. & Esquivel, R. N. Archaeal type IV pili and their involvement in biofilm formation. Front Microbiol 6, 190 (2015).

17 Albers, S.-V. & Jarrell, K. F. The archaellum: an update on the unique archaeal motility structure. Trends Microbiol 26, 351–362 (2018).

18 Moissl, C., Rachel, R., Briegel, A., Engelhardt, H. & Huber, R. The unique structure of archaeal ‘hami’, highly complex cell appendages with nano-grappling hooks. Mol Microbiol 56, 361–370 (2005).

19 Thoma, C. et al. The Mth60 fimbriae of Methanothermobacter thermoautotrophicus are functional adhesins. Environ Microbiol 10, 2785–2795 (2008).

20 Doddema, H. J., Derksen, J. W. & Vogels, G. D. Fimbriae and flagella of methanogenic bacteria. FEMS Microbiol Lett 5, 135–138 (1979).

21 Sarbu, C. Untersuchung der Mth60-Fimbrien von Methanothermobacter thermoautotrophicus Doctoral thesis thesis, University of Regensburg, (2013).

22 Wirth, R., Näther, D. J., Rachel, R., Wanner, G.. Adhesives based on proteins; Adhesives based on derivatives thereof. Germany patent WO2006128678A1 (2006).

23 Fink, C. et al. A shuttle-vector system allows heterologous gene expression in the thermophilic methanogen Methanothermobacter thermautotrophicus ΔH. mBio, e0276621 (2021).

24 Pozzi, R. et al. Distinct mechanisms contribute to immunity in the lantibiotic NAI-107 producer strain Microbispora ATCC PTA-5024. Environ Microbiol 18, 118–132 (2016).

25 Balch, W. E. & Wolfe, R. New approach to the cultivation of methanogenic bacteria: 2-mercaptoethanesulfonic acid (HS-CoM)-dependent growth of Methanobacterium ruminantium in a pressureized atmosphere. Appl. Environ. Microbiol. 32, 781–791 (1976).

26 Sambrook, J., Fritsch, E. F. & Maniatis, T. Molecular Cloning: A Laboratory Manual. (Cold spring harbor laboratory press, 1989).

27 Heap, J. T., Pennington, O. J., Cartman, S. T. & Minton, N. P. A modular system for Clostridium shuttle plasmids. J Microbiol Methods 78, 79–85 (2009).

28 Huff, J. The Airyscan detector from ZEISS: Confocal imaging with improved signal-to-noise ratio and super-resolution. Nat Methods 12, i–ii (2015).

29 Hadley, W. Ggplot2: Elegrant graphics for data analysis. (Springer, 2016).

30 Core Team, R. R: A language and environment for statistical computing. R Foundation for statistical computing, Vienna (2013).

31 Santangelo, T. J. et al. Polarity in archaeal operon transcription in Thermococcus kodakaraensis. J Bacteriol 190, 2244–2248 (2008).

32 Darcy, T. J. et al. Methanobacterium thermoautotrophicum RNA polymerase and transcription in vitro. J Bacteriol 181, 4424–4429 (1999).

33 Metcalf, W. W., Zhang, J. K., Apolinario, E., Sowers, K. R. & Wolfe, R. S. A genetic system for archaea of the genus Methanosarcina: Liposome-mediated transformation and construction of shuttle vectors. Proc Natl Acad Sci U S A 94, 2626–2631 (1997).

34 San Millan, A. et al. Small-plasmid-mediated antibiotic resistance is enhanced by increases in plasmid copy number and bacterial fitness. Antimicrob Agents Chemother 59, 3335–3341 (2015).

35 Majernik, A. I., Lundgren, M., McDermott, P., Bernander, R. & Chong, J. P. DNA content and nucleoid distribution in Methanothermobacter thermautotrophicus. J Bacteriol 187, 1856–1858 (2005).

36 Hildenbrand, C., Stock, T., Lange, C., Rother, M. & Soppa, J. Genome copy numbers and gene conversion in methanogenic archaea. J Bacteriol 193, 734–743 (2011).

37 Zeikus, J. & Wolfe, R. Fine structure of Methanobacterium thermoautotrophicum: Effect of growth temperature on morphology and ultrastructure. J Bacteriol 113, 461–467 (1973).

38 Yoshinaga, M. Y. et al. Methanothermobacter thermautotrophicus modulates its membrane lipids in response to hydrogen and nutrient availability. Front Microbiol 6, 5 (2015).

39 Pfeifer, K. et al. Archaea biotechnology. Biotechnol Adv, 107668 (2021).

