## Supplementary information for "The targeted deletion of genes responsible for expression of the Mth60 fimbriae leads to loss of cell-cell connections in *M. thermautotrophicus* ΔH"

### S1. Supplementary Material and Methods

#### Primer, gBlock, and plasmid lists (Supplementary Table S1-S3)

**Supplementary Table S1**. Primer list

| Name | Purpose | Sequence | Reference |
|---|---|---|---|
| **Seq_CF1** | Analysis genome integration Δ*mth*58-61 | CCACCAGTTCGACTCCCTGG | Fink*, et al.* [1] |
| **Seq_CF2** | Analysis genome integration Δ*mth*58-61 | CTGTTAAAGGCGGGGGTGG | Fink*, et al.* [1] |
| **Seq_CF3** | Analysis genome integration Δ*mth*58-61 | CTTGGGTGATGATGGGATGTATTG | Fink*, et al.* [1] |
| **Seq_CF4** | Analysis genome integration Δ*mth*58-61 | CGAGGAGAAACACATCCAGCTG | Fink*, et al.* [1] |
| **Seq_CF5** | Analysis pMVS1111A:P$_{hmtB}$-*mth*58-61 | GTTAATCCAGCACATCCTCC | Fink*, et al.* [1] |

| | | | |
|---|---|---|---|
| **Seq_CF6** | Analysis pMVS1111A:$P_{hmtB}$-*mth*58-61 | CCTGTCCAACTTATACCTTTGG | Fink*, et al.* [1] |
| **Gib_CF15** | Cloning pCF702 | CGGCTCTAGCTATGTCCGATC | Fink*, et al.* [1] |
| **M13 FW** | Cloning pCF702 | TGTAAAACGACGGCCAGT | Eurofins genomics (Konstanz, Germany) standard primer list |
| **Res_CF3** | Cloning pCF702 | TAGCCTATTCGGCGCGCCGAAATCCCAACCTTCATATAATATTGCAAG | this study |
| **Res_CF4** | Cloning pCF702 | TCATGGCTACGTCGACCGGGCCATAACACATACCAC | this study |
| **Res_CF5** | Cloning pCF702 | TCAGGATTCGGGCCGGCCTCCGTAAGAAGGGAATTGAACCTCC | this study |
| **Res_CF6** | Cloning pCF702 | CGTTGCAATGCATATGGGTACTGTGTGAGGGTCATATTCTG | this study |
| **Res_CF7** | Cloning pCF702 | TATTGCAATGGTCGACAGTGGGCAAGTTGAAAAATTCAC | this study |
| **Res_CF8** | Cloning pMVS1111A: $P_{hmtB}$-*mth*58-61 | ATGCCATTACGGCGCGCCTCACCACACTGTGGCTCCG | this study |
| **Res_CF9** | Cloning pCF702 | TACGTTGCCAGGCCGGCCCCCATGAACCAACCGATGGC | This study |
| **Res_CF10** | Cloning pMVS1111A: $P_{hmtB}$-*mth*58-61 | CGTTGATCCGTTAATTAAGTCCAGGAATAAGGAAGACATCCCG | this study |
| **Seq_CF9** | Analysis genome integration Δ*mth*58-61 | GTGGTCCGCTGGCAGATATG | this study |
| **Seq_CF10** | Analysis genome integration Δ*mth*58-61 | GCTGCTTCCCTGAGGATGTC | this study |
| **Seq_CF11** | Analysis genome integration Δ*mth*58-61 | CTGCTCCCTGGGATGCCTTG | this study |
| **Seq_CF12** | Analysis genome integration Δ*mth*58-61 | GTAATCCCGCTTATGGTTGCCC | this study |

**Supplementary Table S2**. gBlock list

| Name | Purpose | Sequence | Reference |
|---|---|---|---|
| **gBlock_ $P_{hmtB}$_P*acI_mth61*** | Promoter exchange from native to $P_{hmtB}$ | CGGCTCTAGCTATGTCCGATCAATCTTAATTAAGCCTGGAGGAATGCCCCCATGAACC AACCGATGGCTCAGAAAAACCTTAAAATTAGCGATATATTTATATAGGATTATATGAA TAGATAATATCACATAAAATGAGGTGGTTAATTATGAGGACGACAGTTATTTCCGTGA TTTTATTGTTTCTAATC | This study |

**Supplementary Table S3**. Plasmid list

| Name | Function | Reference |
|---|---|---|
| **pMTL83151** | Shuttle vector for *Clostridia* spp. | Heap et al. 2009 |
| **pSB1** | Exchange of $P_{mcrB(M.v.)}$ promoter to $P_{synth}$ for $Neo^R$ | Fink et al 2021 |
| **pCF204** | pUC57 vector including $Neo^R$ controlled by $P_{mcrB(M.v.)}$ | Fink et al 2021 |

| | | |
|---|---|---|
| **pMSV1111A:P$_{hmtB}$-*bgaB*** | Shuttle vector construct including β-galactosidase (*bgaB*)-gene and promoter P$_{hmtB}$ | Fink et al 2021 |
| **pCF702** | flanking regions of *mth58-61* with Neomycin resistance in between controled by P$_{synth}$ | this study |
| **pMVS1111A:PhmtB-*mth58-61*** | *mth58-mth61* operon from *M. thermautotrophicus* ΔH controlled by P$_{hmtB}$ promoter | this study |

### Detailed 3-step cloning strategy for a suicide-vector construct in *M. thermautotrophicus*

The plasmid pCF204, which contains the pUC57 vector backbone and Neo$^r$ with P$_{mcrB(M.v.)}$, flanked by the restriction enzyme recognition sites *Fse*I and *Asc*I, was used as a starting point for the three-step cloning strategy. To exchange the vector backbone of pCF204 towards the *E. coli* vector backbone from pMTL83151 [2], including Cam$^r$, ColE1, and the *tra* minigene with Ori-T, the pMTL83151 backbone was PCR amplified to implement a *Sal*I restriction site and PCR purified. The resulting PCR product of pMTL83151 and pCF204 were restriction digested with *Kpn*I and *Sal*I, PCR purified, and ligated with T4 ligase. After confirmation of the backbone exchange, the up- (0.8 kb) and downstream (1 kb) flanking regions were PCR amplified using Q5 hot start high-fidelity polymerase with primer combinations including the respective restriction site. The PCR products were PCR purified. The up- and downstream flanking region fragments were digested with the restriction enzymes *Asc*I and *Sal*I (upstream) and *Fse*I and *Nde*I (downstream), respectively. The fragments were subsequently ligated into the backbone exchanged construct and digested with the corresponding restriction enzymes. The downstream flanking region was implemented first, followed by the upstream flanking region in the third step after confirmation of downstream flanking region implementation. After fusion of the suicide-vector backbone, including the up- and downstream homologous flanking regions, we exchanged the Neo$^r$ with the non-functional P$_{mcrB(M.v.)}$ spacer to selectable marker Neo$^r$ with P$_{synth}$ from pSB1. This was performed *via* restriction/ligation cloning with the modular restriction sites *Fse*I and *Asc*I also facilitating the exchange of selectable markers for *M. thermautotrophicus* ΔH as a module in the pMVS design [1]. The exchanges resulted in the final suicide-vector constructs pCF702 (**Supplementary Table S3**).

### Oxford Nanopore Sequencing

High molecular-weight genomic DNA from the respective *M. thermautotrophicus* ΔH strains was extracted. For further concentration, size exclusion, and higher purification, the genomic DNA AMPure XP (Beckmann Coulter, Brea (CA), USA) magnetic bead clean-up was performed. The ratio of magnetic beads to genomic DNA volume was 2:1, which should exclude most of genomic DNA fragments smaller than 3 kb. The protocol was performed as described in the user manual for PacBio library preparation (PacBio template preparation and sequencing). Genomic DNA was eluted in 15 µL $H_2O_{millipore}$ and stored at 6°C.

We chose the rapid barcoding Kit (Oxford Nanopore Technologies, Oxford Science Park, UK) for library preparation multiplexing of up to 12 samples for the three different *M. thermautotrophicus* ΔH strains to include all three strains multiplexed in the same flow-cell run. All steps were performed as described in the manufacturer's manual. The run performance was adapted from Esquivel-Elizondo*, et al.* [3].

### Analysis of Oxford Nanopore Sequencing

We performed analyses of the Oxford Nanopore sequencing reads with QIAGEN CLC Genomics Workbench (Qiagen, Hilden, Germany). We generated a workflow where the raw sequences were first trimmed according to **Supplementary Table S4**. Short and low-quality sequencing reads were discarded. Afterwards, the trimmed sequencing reads were aligned to references genomes, exported as genbank format files mapped by the Long Reads (beta) algorithm. Furthermore, we connected the reads tracks to the annotation tracks of the reference sequence genbank files for final analysis of the alignment. Unmapped reads were discarded.

**Supplementary Table S4**. Set-up for trimming Nanopore sequencing reads

| **TRIM READS** | |
|---|---|
| Trim using quality scores | true |
| Quality limit | 0.05 |
| Trim ambiguous nucleotides | true |
| Maximum number of ambiguities | 2 |
| Automatic read-through adapter trimming | false |
| Trim adapter list | |
| Trim homopolymers from 5' | false |
| Trim homopolymers from 3' | false |
| polyA | false |
| polyC | false |
| polyG | true |
| polyT | false |
| Remove 5' terminal nucleotides | false |
| Number of 5' terminal nucleotides | 1 |
| Remove 3' terminal nucleotides | false |

| | |
|---|---|
| Number of 3' terminal nucleotides | 1 |
| Trim to a fixed length | false |
| Maximum length | 150 |
| Trim end | Trim from 3'-end |
| Discard short reads | true |
| Minimum length | 50 |
| Discard long reads | false |
| Maximum length | 1000 |

### S2. Supplementary Results

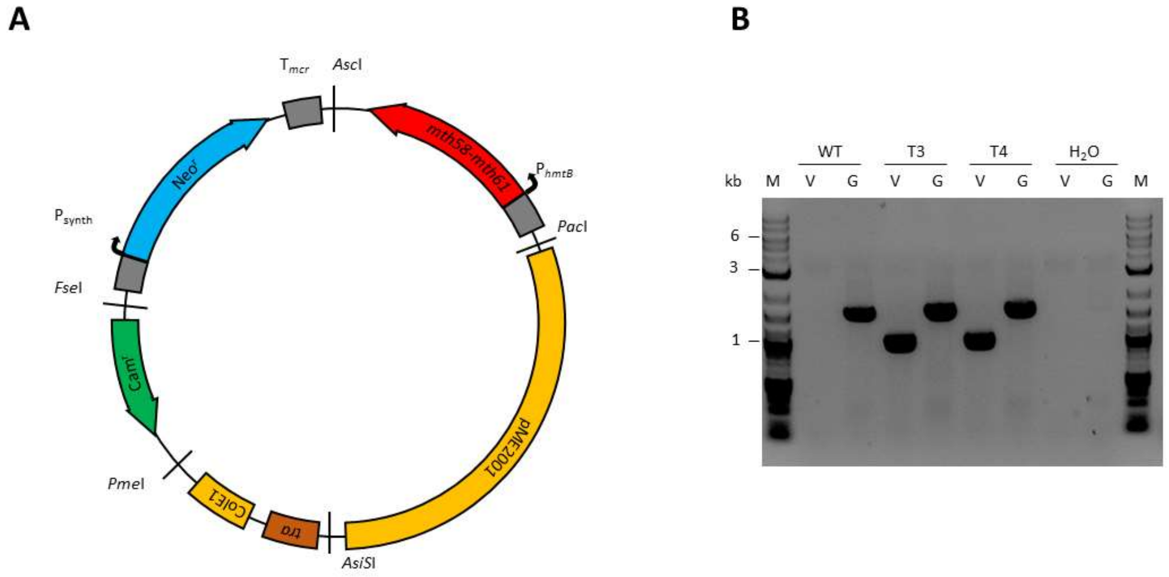


**Supplementary Figure S1**. DNA transfer of shuttle vector construct for constitutive expression of Mth60 fimbriae-encoding operons (**A**) into *M. thermautotrophicus* ΔH confirmed by PCR analysis (**B**). **A)** The pMVS1111A:P$_{hmtB}$-*mth58-61* with neomycin resistance for *M. thermautotrophicus* ΔH (*Neo*$^R$), chloramphenicol resistance for *E. coli* (*Cam*$^R$), origin of replication for *E. coli* (ColE1) including origin of transfer (*tra*), and origin of replication for *M. thermautotrophicus* (pME2001) modules contains the Mth60 fimbriae-encoding operons in the gene of interest module location between *Pac*I and *Asc*I restriction enzyme recognition sites. **B)** The presence of pMVS1111A:P$_{hmtB}$-*mth58-61* in *M. thermautotrophicus* ΔH was confirmed by PCR analysis using the primer combination M (**G**) as control for genomic DNA and Seq_CF5/6 (**V**) for determination of pMVS1111A:P$_{hmtB}$-*mth58-61*. While wild-type *M. thermautotrophicus* ΔH (**WT**) only provides a signal for genomic DNA and the negative control (**$H_2O$**) provides no visible signal, the samples after three (**T3**) and four (**T4**) transfers, respectively, show signals for pMVS1111A:P$_{hmtB}$-*mth58-61* and genomic DNA of *M. thermautotrophicus* ΔH.

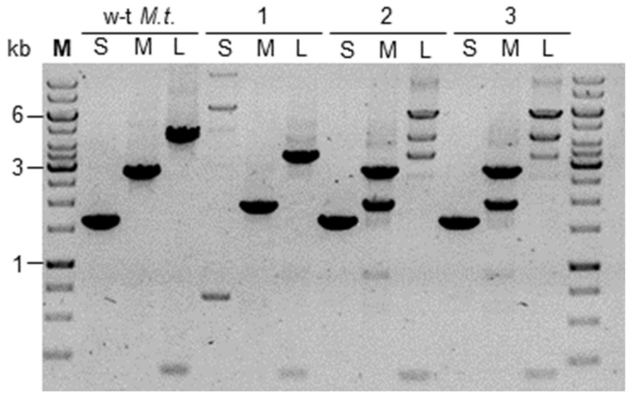


**Supplementary Figure S2**. PCR using S, M, and L primer combination (**main text Figure 1**) of wild-type *M. thermautotrophicus* ΔH (**w-t *M.t.***), a Mth60-fimbriae operon deletion strain (**1**), and two individual clonal populations with mixed signals for wild-type, double-homologous recombination, and single-homologous recombination (**2+3**). Sample 2 was whole genome sequenced using nanopore sequencing.

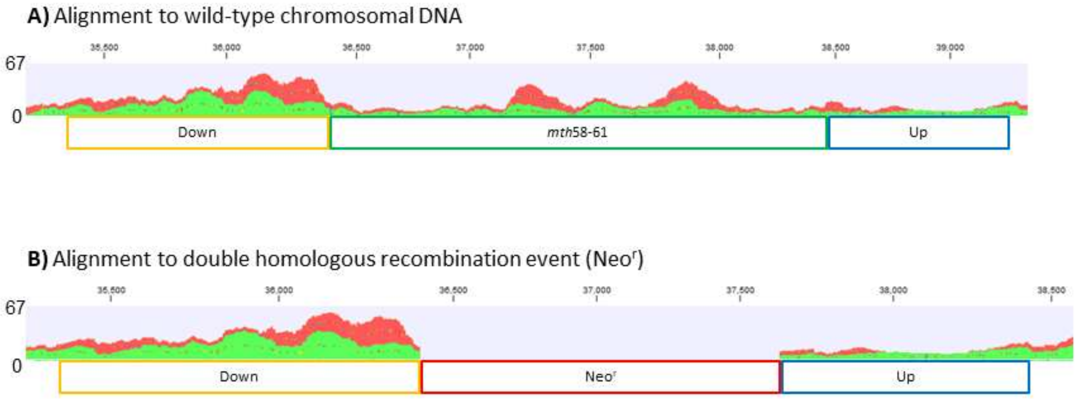


**Supplementary Figure S3**. Sequence alignment of wild-type *M. thermautotrophicus* ΔH nanopore sequencing DNA reads on *in-silico* generated reference genomes of *M. thermautotrophicus* ΔH at the locus of the Mth60 fimbriae-encoding operons and wild-type genomic DNA (**A**) and double-homologous recombined genomic DNA with pCF702 (**B**). The numbers on the Y-axis represent the number of reads extrapolated to 200,000 DNA total reads. Down (**yellow**) represents the downstream homologous region of the Mth60 fimbriae-encoding operons, *mth58-61* (**green**) the Mth60 fimbriae-encoding operons, Neo$^r$ (**red**) the neomycin resistance gene, and up (**blue**) the upstream flanking region of the Mth60 fimbriae-encoding operons.

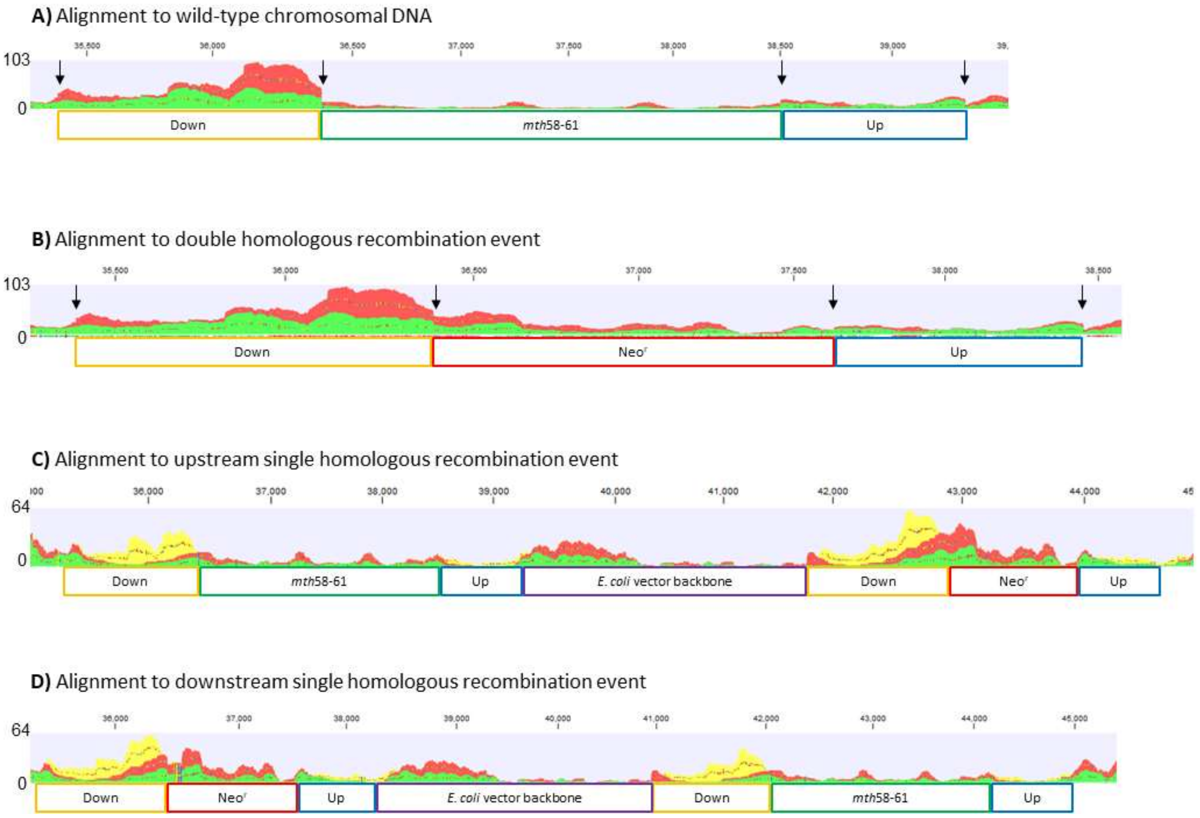


**Supplementary Figure S4**. Sequence alignment of *M. thermautotrophicus* ΔH Δ*mth*60operon:Neo$^r$ nanopore sequencing DNA reads to *in-silico* generated reference genomes of *M. thermautotrophicus* ΔH at the locus of the Mth60 fimbriae-encoding operons; wild-type genomic DNA (**A**), double-homologous recombined genomic DNA with pCF702 (**B**), upstream single-homologous recombined genomic DNA with pCF702 (**C**), and downstream single-homologous recombined genomic DNA with pCF702 (**D**). The **black arrows** in A + B indicate the interface between the homologous flanking region and the Mth60 fimbriae-encoding operons or Neo$^R$ on the inside and to the genomic DNA of *M. thermautotrophicus* ΔH to the outside. The numbers on the Y-axis represent the number of reads extrapolated to 200,000 DNA total reads. Down (**yellow**) represents the downstream homologous region of the Mth60 fimbriae-encoding operons, *mth58-61* (**green**) the Mth60 fimbriae-encoding operons, Neo$^R$ (**red**) the neomycin resistance gene, *E. coli* vector backbone (**purple**), and up (**blue**) the upstream flanking region of the Mth60 fimbriae-encoding operons.

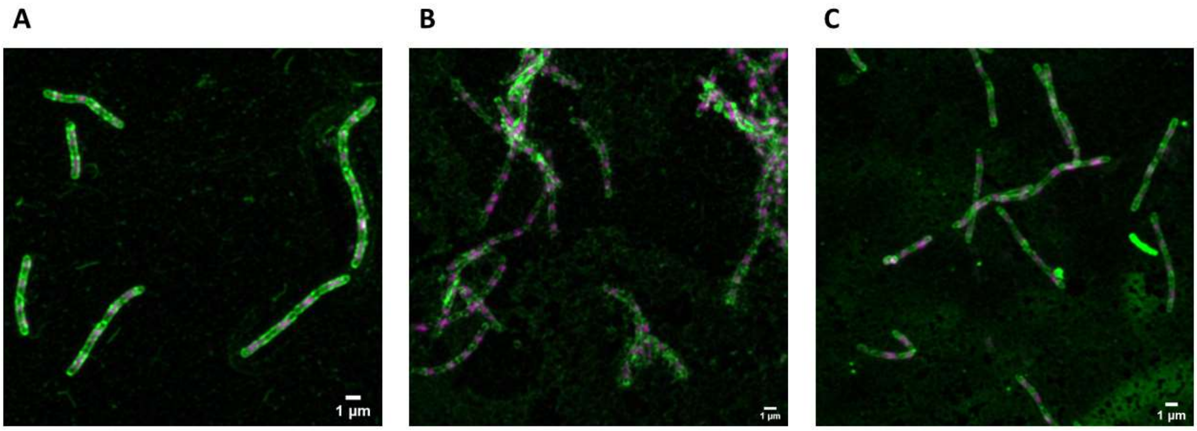


**Supplementary Figure S5.** Larger field of view from **main text Figure 2.** Two-channel maximum intensity z-projection of Airyscan processed z-stacks of immune-fluorescence-stained *M. thermautotrophicus* ΔH strains (A-C). Dapi staining is represented in magenta. The Alexafluor-488 conjugated antibody, which is attached to primary anti-Mth60-fimbriae antibody is depicted in green. **A)** *M. thermautotrophicus* ΔH wild-type. **B)** *M. thermautotrophicus* ΔH containing a shuttle vector for constitutive expression of the Mth60 fimbriae-encoding operons. **C)** *M. thermautotrophicus* ΔH with a deletion of the Mth60 fimbriae-encoding operons.

### Supplementary References


1 Fink, C. *et al.* A shuttle-vector system allows heterologous gene expression in the thermophilic methanogen *Methanothermobacter thermautotrophicus* ΔH. *mBio*, e0276621 (2021).

2 Heap, J. T., Pennington, O. J., Cartman, S. T. & Minton, N. P. A modular system for *Clostridium* shuttle plasmids. *J Microbiol Methods* **78**, 79-85 (2009).

3 Esquivel-Elizondo, S. *et al.* The isolate *Caproiciproducens* sp. 7D4C2 produces n-caproate at mildly acidic conditions from hexoses: genome and rBOX comparison with related strains and chain-elongating bacteria. *Front Microbiol* **11**, 3335 (2021).
